# Concanavalin A as a pan-eukaryotic nuclear envelope marker for expansion microscopy

**DOI:** 10.64898/2026.09.08.750128

**Authors:** Baukje Hoogenberg, Felix Mikus, Priyesh Singh Parihar, Iva Verbanac, Diego Beltrame, Marine Olivetta, Thomas A. Richards, Gautam Dey, Omaya Dudin

## Abstract

Across eukaryotes, the nuclear envelope exhibits distinct remodelling strategies during mitosis: complete breakdown (open mitosis), partial disruption (intermediate), or full retention (closed). However, both light and electron microscopy of the nuclear envelope, at sufficient resolution to determine integrity and mode of remodelling, have proven technically challenging. Here, Concanavalin A (ConA), a glycan-binding plant lectin, provides discernible nuclear envelope/endoplasmic reticulum (NE/ER) labelling in Ultrastructure Expansion Microscopy (U-ExM). Applied post-expansion, ConA circumvents antibody optimisation and labels the NE across mammalian cells and diverse microbial eukaryotes. By imaging four opisthokont and amoebozoan species with established mitotic strategies, *C. perkinsii, D. discoideum, S. pombe*, and *S. arctica*, we distinguished the full spectrum of NE remodelling. Applied to species with poorly characterised mitotic strategies, ConA revealed intermediate mitosis with polar fenestrae in the multinucleate stramenopile *A. limacinum* and a life cycle-coupled switch between open and closed mitosis in the amoebozoan *P. polycephalum*. These observations support the hypothesis that multinucleated life cycles favour closed or intermediate mitosis, protecting chromosomes from capture by neighbouring spindles during synchronous divisions. Together, these results establish ConA in combination with expansion microscopy as a broadly applicable tool for uncovering the diversity and evolution of NE remodelling across eukaryotes.

## Introduction

Compartmentalisation of the genome by a nuclear envelope (NE) that is continuous with the endoplasmic reticulum (ER) is a hallmark of eukaryotic cells, but the composition and architecture of this compartment vary markedly across the eukaryotic tree. Nuclear organisation is highly dynamic, and the NE must be remodelled at every nuclear division. Two broad strategies have evolved across eukaryotes: open mitosis, in which the NE disassembles at mitotic entry and reassembles around segregated chromosomes, and closed mitosis, in which the NE remains intact while the spindle assembles within the nuclear compartment (Dey & Baum, 2021). Between these two extremes lies a continuum of intermediate strategies, including those in which the NE is maintained except for discrete polar fenestrations (Heath, 1980; Sazer et al., 2014). Although open and closed mitosis are well characterised in animal and fungal model systems, the prevalence and distribution of intermediate strategies across the broader eukaryotic tree remain poorly understood. Testing this, however, is limited in part by the tools available to image NE dynamics outside a narrow set of well-studied lineages.

Imaging the NE across diverse eukaryotes comes with several challenges. Lamins and LEM domain proteins, the components routinely labelled in animal nuclei, can be targeted with antibodies but are not conserved throughout eukaryotes. Antibodies against NPC components, including the widely used mAb414, which targets FG repeat nucleoporins, serve as a proxy label across opisthokonts but labelling is often heterogenous or incomplete outside of model species. Lamins, LEM domain proteins and nucleoporins also cycle on and off the NE with cell state and cell cycle stage, so their presence reports the envelope only indirectly. Membrane dyes such as BODIPY TR ceramide label the NE, but also every other membrane in the cell, and standard immunofluorescence lacks the resolution to capture NE features at the nanometre scale (Liffner & Absalon, 2021; Sheard et al., 2023). Most microbial eukaryotes are not genetically tractable, so fluorescent tagging of NE components is not an option. Electron microscopy resolves the ultrastructure but remains low throughput and hard to combine with multi-protein labelling across large species panels. Cryofixation and volume EM have corrected the inconsistent NE preservation of older protocols, yet even in tomograms the NE is not always separable from bulk ER, particularly during mitosis when NPCs may be absent.

Expansion microscopy protocols, such as Ultrastructure Expansion Microscopy (U-ExM), address some of these limitations. U-ExM physically expands the sample fourfold before labelling, improving both spatial resolution and antibody accessibility in a single step (Gambarotto et al., 2019; Mikus et al., 2025; Shah et al., 2024). Accessibility is a critical advantage in microbial eukaryotes, where diverse cell walls often limit antibody penetration in classical protocols. Applied to over 200 species, U-ExM has revealed 3D cytoskeletal and organellar ultrastructure across the eukaryotic tree at a throughput that electron microscopy cannot yet match (Mikus et al., 2025). Antibody-based NE markers, however, are largely restricted to Opisthokonta, while pan-labelling dyes are insufficiently specific (Liffner & Absalon, 2021; M’Saad & Bewersdorf, 2020; Sheard et al., 2023). A broadly compatible NE marker for U-ExM is therefore still lacking.

Concanavalin A (ConA), a lectin from the jack-bean *Canavalia ensiformis* that selectively binds terminal α-mannose and α-glucose residues on glycoproteins, has been widely used to immobilise cells, including yeast, for microscopy and to stain the cell outline in archaea and fungi (Caloca et al., 2022; Cezanne et al., 2023; Tkacz et al., 1971). Beyond these applications, ConA-reactive material has been detected across phylogenetically diverse eukaryotes, from apicomplexan parasites to fish epithelial cells, where it labels glycoproteins along the secretory pathway and the ER/NE network (Bereiter-Hahn, 1990; Luk et al., 2008). More than fifty years ago, Monneron and Segretain showed by electron microscopy that ConA binds extensively to both the inner and outer nuclear membranes of calf thymocytes (Monneron & Segretain, 1974). Given that glycoproteins are broadly conserved across eukaryotes, that ConA binding spans both sides of the nuclear membrane, and that U-ExM overcomes the resolution and accessibility limitations of classical immunofluorescence, we asked whether ConA could serve as a broad-spectrum NE/ER marker across the eukaryotic tree.

We recently showed, using Ichthyosporea, a clade of close animal relatives, that mitosis has diverged towards either a fungal-like closed mode or an animal-like open mode. This divergence may reflect the distinct multinucleated and uninucleated life cycles found across the clade. These findings suggested that multinucleated life cycles favour the evolution of closed mitosis (Shah et al., 2024). However, this hypothesis was based on a limited set of opisthokont species in which NE dynamics could be visualised using tractable markers. Many lineages in which multinucleated or coenocytic life cycles are common were not included. Testing the hypothesis across a broader phylogenetic range could determine whether this constraint is deeply rooted in eukaryotic history. Alternatively, it could reveal that closed and intermediate forms of mitosis have evolved repeatedly in association with multinucleation. Here, we establish ConA as a broad NE/ER marker for U-ExM in fifteen species spanning six eukaryotic supergroups, and show that it distinguishes open, intermediate and closed mitosis.

## Results

While screening commercially available fluorescent probes for U-ExM, we found that conjugated ConA efficiently labels the NE and ER in cultured mammalian cells (Fig. 1B). ConA has previously been used as a general membrane marker in permeabilised, non-expanded cells (Zaganelli et al., 2025). Consistent with this, we observed internal membrane labelling under these conditions in RPE-1 cells (Fig. S1A). In non-permeabilised cells, ConA signal was largely restricted to the cell surface, consistent with lectin binding to plasma membrane glycoproteins (Fig. S1A).

**Fig 1.**
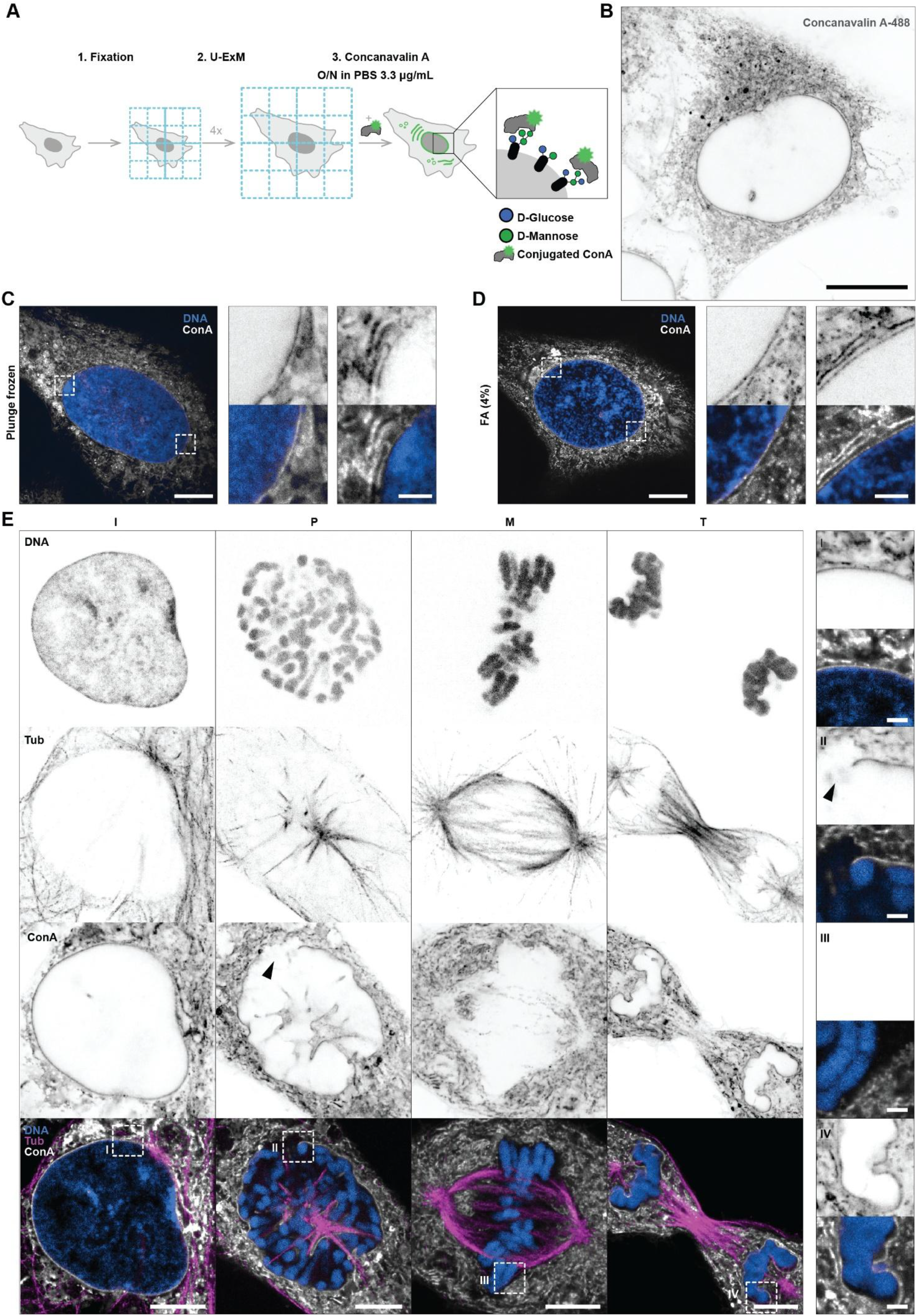
Concanavalin A labels the nuclear envelope in mammalian cells. A. Schematic of the Concanavalin A labelling approach B. Representative expanded RPE-1 cell stained with ConA. A maximum intensity projection (MIP) over two central confocal slices is shown. C. & D. Comparison of plunge frozen (vitrified, C) and chemically fixed (4% formaldehyde [FA], D) RPE-1 cells labelled with ConA (grey) and Hoechst (blue) shown as MIPs over central nuclear regions. Close ups, representing single confocal slices, are indicated by dashed boxes. E. Mitotic stages (Interphase (I), prophase (P), metaphase (M), and telophase (T)) of formaldehyde fixed RPE-1 cells are shown, labelled with Hoechst (blue), tubulin (magenta), ConA (grey). Close up of the chromatin/cytoplasm interface representing dashed boxes are shown. All images are single confocal slices if not stated differently. All scale bars are adjusted for the expansion factor and represent 5µm in overview images and 1µm in close ups.

In contrast, post-expansion ConA labelling produced a robust and homogeneous signal predominantly at the NE and ER. The ∼4-fold increase in resolution enabled clear visualisation of stacked membrane structures and putative Golgi stacks (Fig. 1B and Fig. S1B-D). In U-ExM, ConA was applied at 3.3 µg/ml in PBS overnight at room temperature after immunofluorescence and NHS-ester staining (Fig. 1A). Similar labelling was obtained in PBS-Tween-20 (0.2%) or PBS-Tween-20 supplemented with 3% BSA. In contrast, sodium bicarbonate buffer (100 mM, pH 8) was incompatible with ConA labelling (Fig. S1E). We compared cryofixation and chemical fixation to determine whether gold-standard cryopreservation was necessary. In both cryofixed and chemically fixed RPE-1 cells, ConA produced a comparable sharp, continuous signal at the NE and labelled ER sheets and cytoplasmic vesicles (Fig. 1C&D). In the perinuclear region, the ConA signal showed elongated stacked sheets (Fig. 1C, D crops). At 4-fold expansion, the NE was resolved as a single band approximately 300 nm wide (Fig. S1C, D). Although chemical fixation with formaldehyde produced labelling comparable to cryofixation and is considerably simpler to implement (Mikus et al., 2025; Shah et al., 2024), vitrification remains essential for successful expansion in some species and substantially reduces fixation artefacts (Flori et al., 2025; Hinterndorfer et al., 2022; Laporte et al., 2022). The pan-labelling dyes currently used in expansion microscopy did not match this specificity. NHS ester, which labels total protein, and BODIPY TR ceramide, which labels all membranes, both partly marked the NE and ER but resolved neither with the contrast or the sharpness obtained with ConA (Fig. S1F). As further validation, ConA clearly visualised NE breakdown in mitotic RPE-1 cells: interphase cells showed a continuous NE signal surrounding the DNA, whereas metaphase cells showed loss of the nuclear rim signal, with ConA signal restricted to ER sheets surrounding the condensed chromosomes before staining the reforming NE during telophase (Fig. 1E). ER and NE morphology was comparable between RPE-1 and HeLa cell lines in interphase, while mitotic HeLa cells showed a ConA-intense ER cisternal network, as previously shown in live HeLa cells (Lu et al., 2009; Sengupta et al., 2015) (Fig. S1G). Together, these results establish ConA as a specific, practical, and commercially available NE and ER marker for U-ExM in mammalian cells, requiring no antibody incubation.

To determine what ConA labels within the cell, we co-stained FA-fixed RPE-1 cells with an antibody against KDEL, a canonical ER luminal marker. ConA and KDEL signals co-localised across ER sheets and at the NE, confirming that a large fraction of ConA labelling corresponds to the ER/NE network (Fig. 2A). While KDEL labelling was restricted to ER sheets and tubules, ConA labelled additional glycan-rich structures in the cytoplasm. Across the ER/NE network specifically, ConA produced a stronger signal than KDEL (Fig. 2B, C). Because ConA labelling largely overlaps with the ER/NE network, where proteins can carry different glycan modifications, including N- and O-linked glycans, we next sought to determine which glycan species contribute to ConA binding. Although many eukaryotic glycoproteins are modified with N-linked glycans, O-linked mannosylation also occurs in the ER and is conserved across eukaryotes (Apweiler et al., 1999; Neubert & Strahl, 2016).

**Fig 2.**
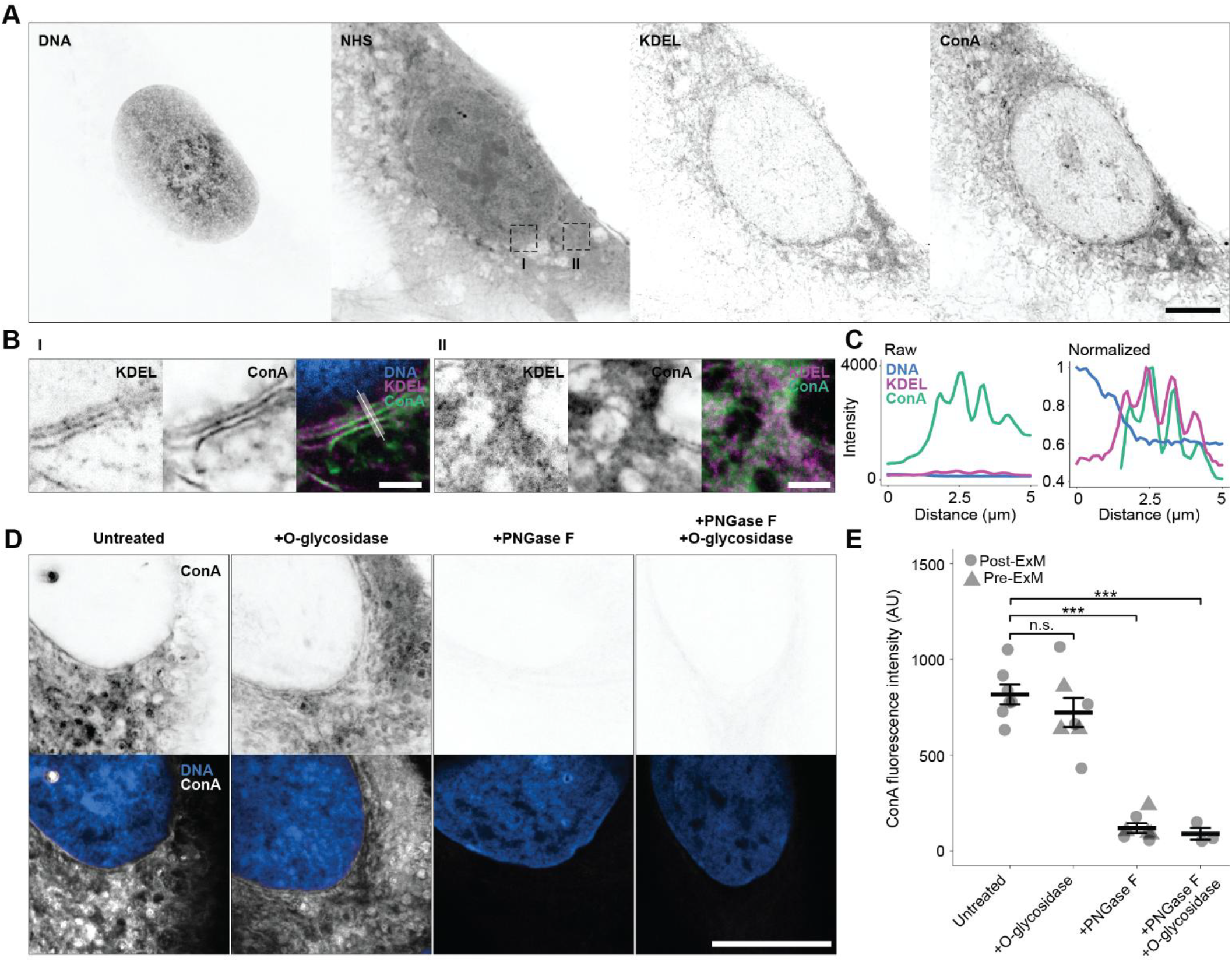
ConA labels the ER by recognising N-linked glycosylation. A. Single channels of an expanded RPE-1 cell co-labelled for DNA, proteome (NHS), the ER (KDEL), and with ConA. A MIP across a central nuclear region is shown. B. Zoom-in of regions indicated in the NHS channel in A showing ER stacks at the nuclear border and cytoplasmic areas. C. Line scan (width 10) across DNA/KDEL/ConA of panel B-I. Raw values and values normalised to the maximum are shown. Distances are adjusted to the expansion factor. D. Cryo-fixed RPE-1 untreated, treated with O-glycosidase, PNGase, or a combination of both post-expansion and stained with Hoechst (blue) and ConA (grey). Channel brightness is scaled the same across all conditions. E. Fluorescence intensity measured for ConA (AU). C/O/N N=7, O+N N=3. N indicates number of cells. Pre-ExM (circle) post ExM (triangle). A one-way ANOVA determined significant effect of treatment on ConA intensity, F (3,20) = 44.66, p<0.001. Dunnett’s test p values for N and O+N conditions were <0.001 (***), for O p = 0.468 (ns). All images are single confocal slices if not stated differently. All scale bars are adjusted for the expansion factor and represent 5µm in overview images and 1µm in close ups.

To identify the glycan species responsible for ConA binding and the linkage through which it is attached, we treated fixed cells with deglycosylation enzymes PNGase F, which cleaves N-glycans, O-glycosidase, which removes O-glycans, or both enzymes in combination, either before or after expansion (Fig. 2D, E). Treatment with O-glycosidase alone did not affect ConA signal intensity compared to untreated controls (Fig. 2E) while PNGase F treatment caused a strong, significant reduction in ConA fluorescence intensity (p < 0.001; Fig. 2D, E).

At high contrast settings, a residual ConA signal remained after digestion with PNGase F, reflecting a potential minor contribution of non-N-linked glycosylation or incomplete enzymatic digestion. Taken together, these results show that ConA binds primarily to N-linked glycans in the ER/NE network, consistent with results obtained previously by electron microscopy (Bereiter-Hahn, 1990; Monneron & Segretain, 1974).

To assess whether ConA could be used to label the NE beyond mammalian cells, we applied the same U-ExM protocol across a panel of 15 species spanning six eukaryotic supergroups (Fig. 3A, S2). The panel comprised five opisthokonts (*Chromosphaera perkinsii, Sphaeroforma arctica, Schizosaccharomyces pombe, Schizosaccharomyces japonicus* and *Choanoeca flexa*), two amoebozoans (*Dictyostelium discoideum* and *Physarum polycephalum*), four SAR species (*Aurantiochytrium limacinum, Toxoplasma gondii, Euplotes rariseta* and a Ross Sea dinoflagellate (RSD)), two discobids (*Trypanosoma brucei* and *Naegleria gruberi*), one metamonad (*Giardia lamblia*) and one archaeplastidan (*Chlamydomonas reinhardtii*). In most species, ConA labelling produced a clear and continuous signal at the NE together with a resolved ER network (Fig. S2B-E, S2H and S2J), showing that ConA-reactive N-linked glycans at the NE/ER are broadly distributed across the eukaryotic tree. Labelling was fainter in *Giardia lamblia* and *Choanoeca flexa*, where ConA outlined the nucleus but did not resolve a distinctive ER network (Fig. S2G, I). A complete lack of signal was observed in the parasite *Toxoplasma gondii*, although DNA and tubulin labelled normally in the same cells (Fig. S2F). *T. gondii* which is known to have a reduced glycosylation machinery and may therefore not produce ConA-reactive glycans across all life stages (Bandini et al., 2019; Lombard, 2016; Tomita et al., 2017). ConA therefore labels the NE and ER in every supergroup sampled, a breadth that no antibody-based NE marker currently matches.

**Fig 3.**
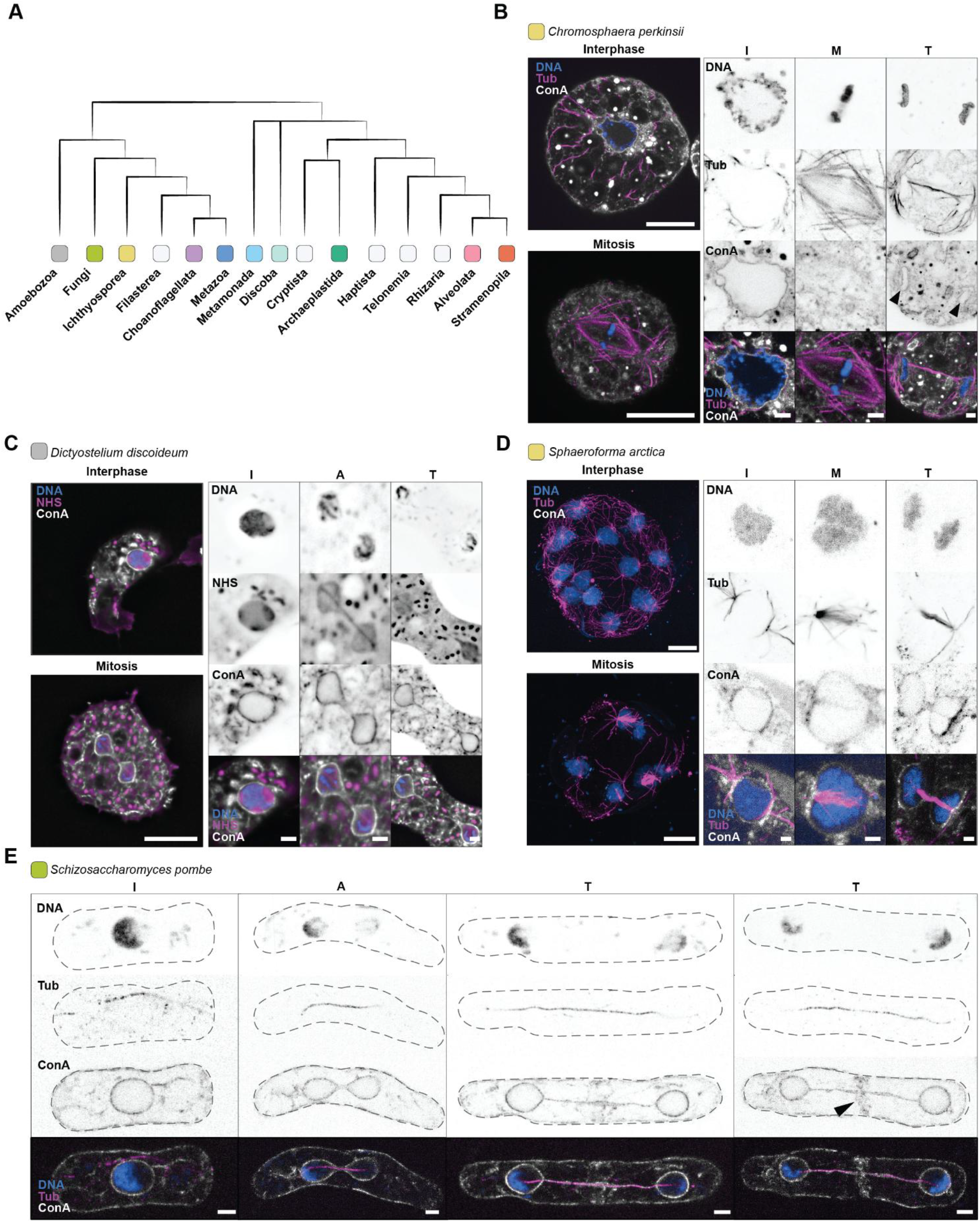
ConA as a readout for open/closed mitosis across diverse eukaryotes. A. Cladogram of eukaryotes with colours indicating the position of used species within present figures. Selected species represent a gradient of open to closed mitotic strategies where known. B. Unicellular mitotic stages of the ichthyosporean *Chromosphaera perkinsii*, FA fixed and labelled with ConA (grey), DNA (blue), and tubulin (magenta). Whole cell images for interphase and metaphase and nuclear crops for interphase (I), metaphase (M), and telophase (T) are shown. Loss of ConA signal surrounding the metaphase chromatin indicates open mitosis with NE reformation during telophase. Arrowheads indicate position of separating nuclei. C. The amoebozoan *Dictyostelium discoideum* labelled with ConA (grey), Hoechst (blue), and NHS ester (magenta). Whole cell images for interphase and metaphase and nuclear crops for interphase (I), anaphase (A), and telophase (T) are shown. ConA signals are retained throughout division with loss occurring only in the spindle midzone. Cells were plunge frozen and MIP projections of central deconvolved widefield slices are shown. D. Coenocytic mitotic stages of the ichthyosporean *Sphaeroforma arctica*, FA fixed and labelled with ConA (grey), DNA (blue), and tubulin (magenta). MIP of whole cells during interphase and metaphase, as well as single confocal slices for nuclear crops of interphase (I), metaphase (M), and telophase (T) cells are shown. ConA signal is maintained around the chromatin throughout mitosis indicating a closed mitosis. E. High pressure frozen (HPF) fission yeast *Schizosaccharomyces pombe* labelled with ConA (grey), Hoechst (blue), and tubulin (magenta). Whole cell images for interphase (I), anaphase (A), and telophase (T) are shown. Arrowhead indicates ConA labelling at the septum. All images are single confocal slices if not stated differently. All scale bars are adjusted for the expansion factor and represent 5µm in overview images and 1µm in close ups.

We then evaluated whether NE/ER labelling with ConA could distinguish the modes of nuclear division, from fully open to closed mitosis (Fig. 3B to E). Four opisthokont and amoebozoan species with distinct and well-characterised mitotic strategies were expanded and labelled with ConA. In *C. perkinsii*, ConA labelled the NE during interphase (Fig. 3B). This ConA signal was completely lost during metaphase and reappeared around separating nuclei during telophase, consistent with the open mitosis characterised in this species (Shah et al., 2024).

In *D. discoideum*, ConA labelled the NE and membrane extensions between the separating nuclei (Fig. 3C). The NE remained largely intact throughout mitosis, consistent with the previously described intermediate mode of division in *D. discoideum* (McIntosh et al., 1985). Similarly, in *S. pombe*, the NE remained intact during division, and ConA effectively labelled the mitotic midzone, membrane extensions, and the site of abscission (Fig. 3E). In *S. arctica*, ConA outlined the nucleus continuously throughout division, reflecting the closed mitosis of this species (Fig. 3D) (Shah et al., 2024). Together, these results show that ConA labels the NE/ER across diverse eukaryotes and visualises different mitotic strategies, from complete NE breakdown to fully closed division.

As ConA showed promise for revealing diverse mitotic strategies, we next applied ConA U-ExM to two species for which immunostaining approaches have not assessed the mode of nuclear division at high resolution: the marine thraustochytrid *Aurantiochytrium limacinum* and the slime mould *Physarum polycephalum* (Fig. 4A). Multinucleated coenocytic stages with synchronous nuclear division have been previously reported in the marine thraustochytrid *A. limacinum*, although its mitotic strategy remains poorly characterised (Dellero et al., 2018). In interphase cells, ConA continuously labelled the NE around each nucleus (Fig. 4A). As cells entered prophase and early spindles were formed, the NE remained largely intact (Fig. 4A). At metaphase, the NE was maintained except for discrete fenestrations at each spindle pole, visible as clear gaps in the ConA signal flanking the centrioles identified by tubulin staining (Fig. 4A, crops). NE/ER remnants persisted at both ends of the metaphase plate, surrounding the condensed chromosomes (Fig. 4A). Thus, *A. limacinum* does not undergo the extensive NE breakdown characteristic of open mitosis. Instead, this pattern, characterised by a largely intact NE with polar fenestrations accommodating the centriolar spindle poles, is consistent with an intermediate mitotic strategy. A comparable pattern has previously been reported by electron microscopy in the related thraustochytrid *Thraustochytrium* sp. (Kazama, 1974). Coenocytic growth and a retained NE therefore co-occur outside Obazoa (Opisthokonta, Apusomonadida and Breviatea). Either the coupling between coenocytic growth and a retained envelope dates to the common ancestor of amorpheans and stramenopiles, close to the root of the eukaryotic tree, or it has been assembled anew in every lineage that became coenocytic.

**Fig 4.**
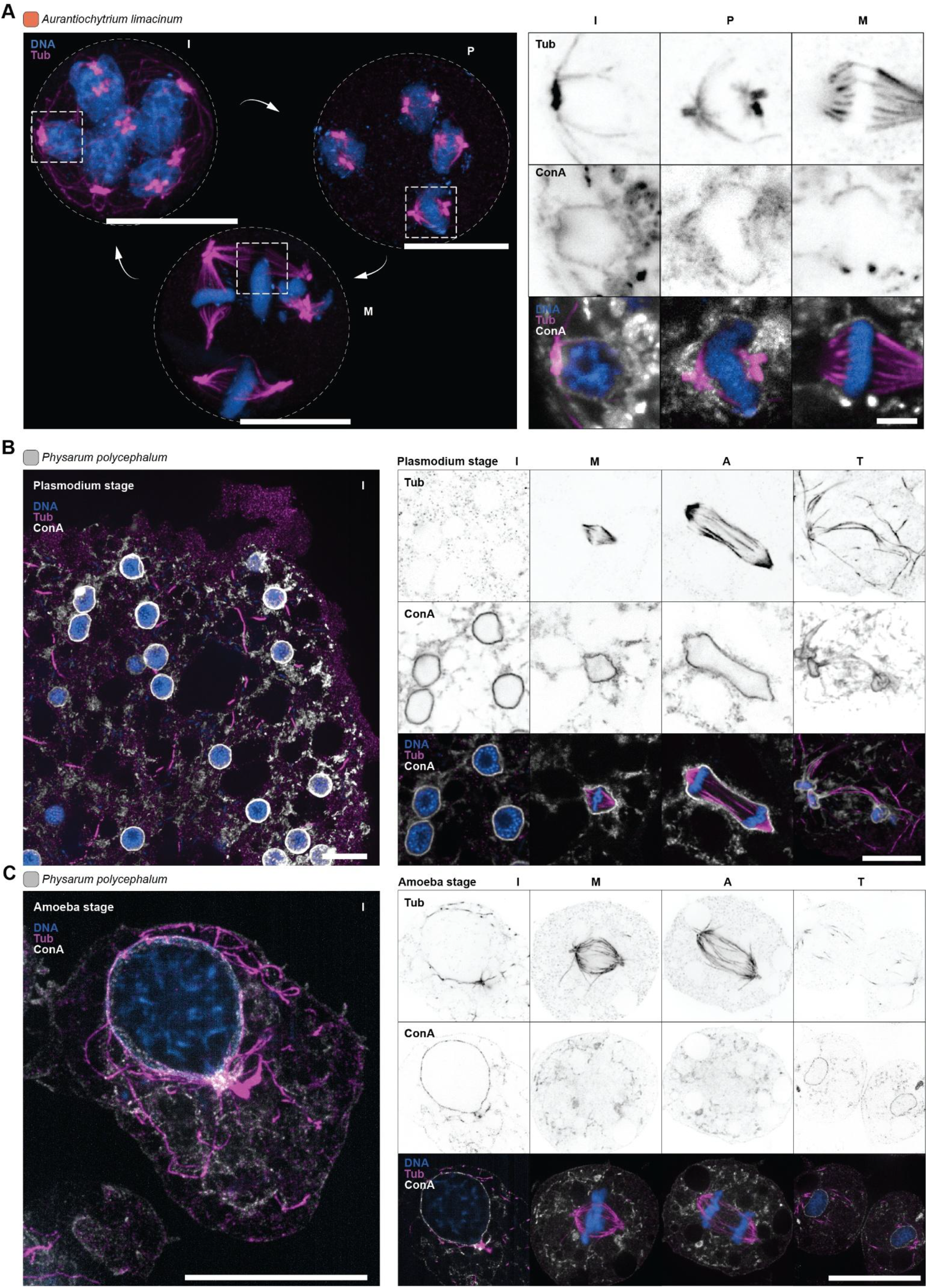
ConA reveals mitotic strategy in the thraustochytrid *A. limacinum* and strategy switching across life stages in the slime mould *P. polycephalum*. A. Multinucleated mitotic stages of *Aurantiochytrium limacinum*, FA fixed and labelled with ConA (grey) and for DNA (blue), and tubulin (magenta). MIP of tubulin and DNA of whole coenocytic cells during interphase (I), prophase (P), and metaphase (M). Single confocal slices of representative nuclei are shown, with ConA identifying polar fenestrations of the NE during pro- and metaphase. B. Overview of *Physarum polycephalum* interphase plasmodial stage. MIP of central confocal slices are shown. Interphase (I), metaphase (M), anaphase (A), and telophase (T) of plasmodial *Physarum polycephalum* labelled with ConA (grey) and for DNA (blue), and tubulin (magenta). ConA signal is retained around the DNA. A MIP of central confocal slices is shown for telophase. C. Overview of *Physarum polycephalum* interphase amoebal stage. MIP of central confocal slices are shown. MIP images of interphase (I), metaphase (M), anaphase (A), and telophase (T) of amoebal *Physarum polycephalum* stages labelled with ConA (grey) and for DNA (blue), and tubulin (magenta). All images are single confocal slices if not stated differently. All scale bars are adjusted for the expansion factor and represent 5µm in overview images and 1µm in close ups.

We next asked whether ConA could be applied to a species with an even starker contrast in mitotic strategy: *P. polycephalum* switches between open mitosis in the uninucleate amoeba and closed mitosis in the multinucleated coenocytic plasmodium (Aldrich, 1969; Burland et al., 1993; Solnica-Krezel et al., 1991). In the plasmodium, ConA outlined each nucleus as a continuous ring throughout division (Fig. 4B). The NE remained intact during metaphase and anaphase, and at telophase nuclei adopted a dumbbell shape, characteristic of closed mitosis, with ConA signal maintained across the constriction (Fig. 4B). In the amoeba, ConA outlined the nucleus in interphase, and this signal was completely lost at metaphase, with no detectable NE signal surrounding the condensed chromosomes (Fig. 4C), consistent with complete NE breakdown. While the life cycle-coupled mitotic switch in *P. polycephalum* has been previously reported (Solnica-Krezel et al., 1991), ConA labelling combined with U-ExM recapitulates both modes in 3D with a single probe, without species-specific antibodies and across developmental stages of one organism. Together, these results establish ConA combined with U-ExM as a route to comparative analyses of how life cycle, ecology and evolutionary history shape remodelling of the nuclear compartment during cell division. A single probe distinguishes open, intermediate and closed mitosis in distantly related lineages.

## Discussion

Imaging the NE across eukaryotic diversity has long been constrained by technical limitations. Antibodies against NE components cross-react poorly outside commonly used model systems, membrane dyes label all membranes indiscriminately, and electron microscopy, while providing high resolution, is expensive and low throughput. We show that ConA addresses many of these limitations. This jack-bean lectin binds N-linked glycans, which are present on membranes across the eukaryotic tree, generating sharp NE/ER signal within the standard U-ExM workflow across fixation conditions. ConA is commercially available, requires no antibody optimisation, is compatible with any fluorophore conjugate, and costs a fraction of what antibody development does.

However, we note some organisms, including *Toxoplasma gondii*, do not stain with ConA. Rather than representing a technical failure, this absence is biologically informative: *Toxoplasma* has undergone genome reduction and streamlined its glycosylation machinery, particularly the lumenal glycosyltransferases that elaborate N-glycan complexity (Luk et al., 2008). The parasite retains functional N-glycosylation, but its simplified and specialised repertoire appears insufficient for ConA recognition, at least during the specific life cycle stage imaged here. *Giardia lamblia* is a different case. ConA labelled the cell and outlined both nuclei, but no distinct ER network was resolved (Fig. S2G). In *Giardia*, the ER is contiguous with the endosomal and lysosomal compartment rather than a discrete organelle (Abodeely et al., 2009). There may therefore be no separate NE/ER network to label. Whether this changes between trophozoites and cysts remains to be tested. More broadly, variation in ConA reactivity across species may reflect differences in the completeness and complexity of each organism’s N-glycosylation machinery, making the presence or absence of signal itself a useful glycobiological readout.

*A. limacinum* is the clearest new case. Its nuclei divide in a shared cytoplasm, and the NE is retained except at the spindle poles. This aligns with our earlier hypothesis that multinucleated life cycles favour closed or intermediate mitosis. It also matches what ConA shows in species whose strategies were already established: open mitosis in *C. perkinsii*, intermediate mitosis in *D. discoideum* and *S. pombe*, and closed mitosis in *S. arctica*. In multinucleated cells undergoing synchronised nuclear divisions, chromosomes exposed to the shared cytoplasm risk capture by microtubules from neighbouring nuclei, and missegregation follows. Maintaining the NE, or breaching it only at discrete and controlled points, removes that risk. Whether the same holds across other stramenopiles, or across SAR, remains to be seen. *Physarum polycephalum* makes the same point within a single genome. The uninucleate amoeba divides with an open NE, the multinucleated plasmodium with a closed one. This life cycle-coupled transition is not a fixed evolutionary difference, but rather a plastic developmental change, suggesting that the transcriptional and regulatory machinery governing mitotic mode is conserved and can be toggled between states. Comparing gene expression between the amoeboid and plasmodial stages, as well as between *A. limacinum* and its uninucleated relatives, may identify the determinants that specify each mode of mitosis. In this way, these developmental and life cycle transitions provide an opportunity to study the mechanistic basis of mitotic evolution.

To understand the distribution and evolutionary origins of mitotic strategies across the eukaryotic tree, we aim to systematically screen microbial eukaryotes spanning major lineages. ConA with U-ExM enables rapid phenotyping at a throughput inaccessible to electron microscopy, allowing us to stage cells during NE breakdown or reformation across hundreds of species. Used as a discovery platform, this approach can guide targeted mechanistic work, including electron microscopy, transcriptomics and genome analysis to relate NE strategy to gene repertoire. This positions ConA with U-ExM to map mitotic evolution and test whether life cycle strategy, ecological context and phylogenetic position predict the mode of NE remodelling.

## Materials and Methods

### Culturing

*Chromosphaera perkinsii* was cultured at 26°C in *C. perkinsii* medium (CpM) containing NaCl (20 g/L), glucose (10 g/L), peptone (5 g/L), yeast extract (3 g/L) and malt extract (3 g/L). *Sphaeroforma arctica* and *Aurantiochytrium limacinum* were cultured in Marine Broth 2216 (Millipore) at 17°C.

*Physarum polycephalum* was cultured as myxamoebae and microplasmodia in semi-defined medium (SDM) at 25°C and 150 rpm. SDM contained glucose (10 g/L), soytone (10 g/L), KH_2_PO_4_ (2 g/L), CaCl_2_·2H_2_O (0.92 g/L), MgSO_4_·7H_2_O (0.6 g/L), FeCl_2_·4H_2_O (0.039 g/L), ZnSO_4_·7H_2_O (0.034 g/L), citric acid (3.54 g/L), EDTA disodium salt (0.224 g/L), biotin (0.005 g/L) and thiamine hydrochloride (0.04 g/L) in distilled water, adjusted to pH 4.6 and autoclaved. Before use, the medium was supplemented with penicillin-streptomycin (1:100) and hemin (1:200 of a 0.05% hematin solution in 1% NaOH).

*Choanoeca flexa* was cultured in 1% seawater complete medium (SWC) at 30°C. SWC consisted of Tropic Marin Salts (24 g/L), Bacto peptone (5 g/L), yeast extract (3 g/L), 0.3% Glycerol in MQ water. SWC was autoclaved and filtered before use.

All other species were received as fixed samples, either FA-fixed or high pressure frozen, as indicated in the figure legends.

### Chemical and cryofixation

For chemical fixation, cells were fixed in 4% formaldehyde (FA) (Sigma F8775-500ML) for 30 min at room temperature (RT), with or without 0.025% glutaraldehyde (GA) (Sigma G6257-10ML). After three washes in PBS, samples were stored at 4°C. For cryofixation, cells were plated onto poly-D-lysine (Sigma P1024)-coated coverslips and manually plunge-frozen into liquid ethane. The fission yeasts S. *pombe* and *S. japonicus* were concentrated by centrifugation at 1.500 rcf and high pressure frozen using a Leica EM ICE in 6mm planchettes. Cryofixed samples were transferred to cryogenic acetone (Acros Organics 326801000) with 0.5% FA and 0.02% GA chilled by dry ice and allowed to gradually warm to room temperature overnight (ON). Rehydration was performed by stepwise transfer through ethanol solutions (100%, 95%, 75%, 50%, 0%) (ThermoScientific E/0650DF/P17), followed by storage in PBS at 4°C. (Gambarotto et al., 2019; Laporte et al., 2022)

### Ultrastructure Expansion Microscopy (U-ExM)

U-ExM was performed as previously described (Gambarotto et al., 2019). Briefly, samples were anchored ON at 37°C in 0.7% FA and 1% acrylamide (Sigma #A4058) in PBS. Samples were then embedded in a swellable hydrogel consisting of 19% (w/w) sodium acrylate (Combi-Blocks #QC-1489); 10% (w/w) acrylamide (Sigma #A4058); 0.1% (w/w) N,N’-methylenebisacrylamide (Sigma #M1533). Polymerisation was initiated by adding 10% (w/w) ammonium persulfate (APS) (ThermoScientific #17874) and 10% (w/w) TEMED (ThermoScientific #17919) and proceeded for 1 h at 37°C. Gels were denatured for 1 h at 95°C in denaturation buffer (200 mM SDS (CarlRoth #CN30.2), 200 mM NaCl (Sigma #S5886), 50 mM Tris (Biosolve #0020092391BS) pH 9.0), then expanded through multiple water exchanges. For gel mounting, gels were cut to appropriate sizes and attached to precoated poly-D-lysine coverslips and sealed using i-Spacers (Sunjin Lab #IS013). The expansion factor was determined for every gel by comparing gel diameter before and after expansion, and all distances and scale bars reported here are corrected by the factor measured for the gel they were acquired from. Across gels and species, expansion factors were close to 4-fold.

Immunofluorescence labelling was performed by incubating gels in primary antibody solution (1:300 in PBS (AppliChem #A0965,9010) + 0.1% Tween-20 (AppliChem #A4974,500) + 3% BSA) ON at 37°C, followed by secondary antibody (1:500) for 3 h at 37°C. NHS-ester labelling of total protein was achieved with 1:1000 fluorophore-conjugated N-hydroxysuccinimide ester at a stock concentration of 2 mg/mL (Dyomics #647-01) for 1 h at RT. Membrane labelling was performed with 1:1000 BODIPY TR Ceramide (ThermoScientific #D7540) ON at RT, at a stock concentration of 2 mM. DNA staining was performed with Hoechst at 1:1000, either concurrent with NHS labelling or during the final re-expansion step. Primary antibodies used here include rabbit anti-KDEL (ABCD #AW954), rabbit anti-alpha tubulin and guinea pig anti-alpha tubulin (ABCD #AA345), rabbit anti-beta tubulin and guinea pig anti-beta tubulin (ABCD #AA344). Secondary antibodies used include donkey anti-rabbit Alexa488, 568 and 647 (Invitrogen #A32790, #A10042 and #A32795) and goat anti-guinea pig Alexa647 (Invitrogen #A21450

ConA labelling was performed post-immunofluorescence using Concanavalin A (ConA) conjugated to a fluorophore (Biotium #29016, 488). Optimal results were achieved with ConA at 1:300 in PBS, incubated ON at RT.

Non-expanded controls were permeabilised in PBS + 0.1% Triton X-100 (Sigma #X100-100ML) after FA fixation, then processed identically through immunofluorescence and ConA labelling.

For de-glycosylation experiments, samples were pre-gelation treated with PNGase F (1000 U, New England Biolabs #M0704) and O-glycosidase (100 U, NEB #P0917) for 4 h at 37°C in glycobuffer 2 (NEB #B1004). Post-gelation treatment used denatured gel pieces incubated with PNGase F, O-glycosidase, or both for 2 h at 37°C in glycobuffer 2 + 0.5% NP-40, followed by ConA labelling as described above.

### Microscopy

Three complementary imaging systems were used for this study. Widefield epifluorescence imaging was performed on a ZEISS Axio Observer 7 inverted microscope equipped with a motorised XY stage and joystick control, a motorised Z drive with Definite Focus 2 autofocus, a motorised nosepiece, and an automated Sample Finder for rapid sample localisation and focal plane detection. Fluorescence illumination was provided by a ZEISS Viluma 5 LED light source with excitation peaks at 385 nm, 475 nm, 555 nm, and 630 nm, coupled to four single band-pass emission filters (DAPI 405/50 nm, GFP/Alexa-488 525/50 nm, RFP/Alexa-568 609/54 nm, Cy-5/Alexa-647 690/50 nm). Images were acquired using a Hamamatsu Orca Flash 4 USB 3.0 camera and ZEN 3.11 software. Objectives employed included Fluar 5x/0.25, Plan-Apochromat 20x/0.8 air, C-Achroplan 32x/0.85 water, and LD C-Plan-Apochromat 40x/1.1 water, with an auto-immersion module enabling rapid switching between water-immersion objectives.

Confocal imaging of expanded samples was conducted on two spinning disk confocal microscopes. The laboratory system consisted of a Nikon Eclipse Ti2-E inverted microscope equipped with a motorised Z drive (10 nm step size in open loop, 20 nm in closed loop), motorised XY stage with encoders and joystick control, and Perfect Focus autofocus. This microscope was coupled to a Yokogawa CSU-W1 spinning disk confocal unit with a 50 μm pinhole. Excitation was provided by a ZIVA Light Engine with seven independently controlled laser lines (405, 446, 488, 518, 577, 639, 748 nm) with integrated despeckler. Images were captured using an ORCA-Fusion C14440-20P sCMOS camera (2304×2304 pixels, 6.5×6.5 μm pixel size, 80% quantum efficiency) with CoaXPress interface. The facility spinning disk system was based on a Nikon Ti inverted microscope with Perfect Focus autofocus and a Yokogawa CSU-W1 spinning disk confocal unit, equipped with four excitation lasers (405, 488, 561, 640 nm), a back-illuminated sCMOS camera (1200×1200 pixels, 11×11 μm, 95% quantum efficiency), and temperature control capability (17–37°C). Both spinning disk systems employed water-immersion objectives with extended working distances for imaging expanded gels: CFI Plan Apochromat 20x/0.8 WI (WD 0.55 mm), 40x/1.1 WI (WD 0.62 mm), and 60x/1.2 WI (WD 0.29 mm). A high-resolution 100x/1.49 NA oil immersion objective was also employed for high-magnification imaging. Image acquisition and analysis were performed using NIS-Elements, Zen Microscopy Software, and SlideBook 3i.

### Image analysis

Image analysis was performed using ImageJ (v.1.54p), R (v.4.4.2), and RStudio (v.2026.04.0+526). Figures were assembled with Affinity by Canva (v.3.2.2) and Adobe Illustrator (v.29.02). Widefield z-stacks in Fig. 3C (LD C-Plan-Apochromat 40x/1.1 water; voxel size 162 × 162 × 330 nm) were deconvolved in Fiji/ImageJ (v.1.54p) using the GPU-accelerated, FFT-based non-circulant Richardson–Lucy algorithm implemented in clij2-fft (v2.2.0.22; CLIJ ecosystem, clijx-deconvolution update site), with 120 iterations and no regularisation. A single theoretical PSF (Born & Wolf 3D model, PSF Generator, EPFL; NA 1.1, water immersion n = 1.33, emission 610 nm) was applied to all channels (Haase et al., 2020; Kirshner et al., 2013).

### Quantification and statistical analysis

ConA and KDEL signals were measured over a line plot (line width 10 pixels), drawn as indicated in Fig. 2B, and plotted in Fig. 2C, across the NE and ER. Values were normalised to the maximum value. ConA fluorescence intensity was measured on the images in Fig. 2D and plotted in Fig. 2E. Intensity was determined by drawing 3 boxes per cell, averaging values and subtracting background intensity values, with the cell as the unit of replication. Digestions performed before and after expansion were pooled within each treatment. Statistical analysis was performed in RStudio, using a one-way ANOVA followed by Dunnett’s test comparing treatment means against the untreated control (multcomp package). ConA signal across the NE and ER (Fig. S1C, D) was measured over a line plot (width 1 pixel) across the NE or ER in at least 20 measurements, some measurements taken from the same cell.

### Declaration of generative AI in the writing process

Claude Opus 5 (Anthropic) was used by the authors strictly as a copy-editing tool to improve the readability and language quality of the manuscript. ChatGPT-5 (OpenAI) was used by the authors to generate and augment scripts for data visualisation and statistical analysis using Rstudio. Generative AI played no part in the intellectual conception, study design, data collection, or the generation of figures. The authors oversaw the entire editing process, verified all changes against the raw data, and maintain absolute accountability for the scientific integrity of the work.

## Acknowledgements

We thank the Dey and Dudin Lab members for discussions, Hiral Shah and Mylan Ansel for critical reading and comments on the manuscript. We thank Paula Llanos for support to upload all the data to BioImage archive. We are grateful to Angelique Perret and Thierry Soldati (University of Geneva) for *Dictyostelium discoideum* fixed samples; Emile Roberts and Charlotte Aumeier (University of Geneva) for RPE-1 fixed cells; Cesar Bernat and Aurelien Roux (University of Geneva) for HeLa fixed cells; Alex de Mendoza (Queen Mary University of London) for *Aurantiochytrium limacinum* cultures; Mathieu Funk and Carmen Faso (University of Bern) for *Giardia lamblia* fixed samples; Amandine Guerin (University of Geneva) for *Toxoplasma gondii* fixed samples; Sabrina Absalon (Indiana University School of Medicine) and Julius Lukeš (Czech Academy of Sciences, Prague) for *Trypanosoma brucei* fixed samples; Pierre Gönczy (EPFL) and Alexander Woglar (University of Geneva) for *Naegleria gruberi* fixed samples; Sophie Martin and Laura Merlini (University of Geneva) for *Schizosaccharomyces pombe* and *Schizosaccharomyces japonicus* cultures; Johan Decelle and Ananya Rao (LPCV, CNRS, Grenoble) for Ross Sea dinoflagellate fixed samples; and Caroline Simon (EMBL, Heidelberg) for *Chlamydomonas reinhardtii* cultures. We thank Nicolas Chiaruttini (University of Geneva) for image analysis support regarding deconvolution. We acknowledge support from the University of Geneva Photonic Bioimaging Facility. This work was supported by Swiss National Science Foundation Starting grant no. TMSGI3_218007 (O.D.) and UNIGE core funding; the European Union through an ERC grant (KaryodynEvo, 101078291) to G.D.; the EMBL Planetary Biology Transversal Theme seed grant (G.D. and O.D.); EMBL Core funding to G.D.; and the Gordon and Betty Moore Foundation (GBMF13113) to O.D. and G.D.

## Data availability

All image data related to this study are deposited at BioImage Archive (Hartley et al., 2022) at https://www.ebi.ac.uk/biostudies/bioimages/studies/S-BIAD4075?key=93fe4073-72ce-4b08-b642-00afba063556 (10.6019/S-BIAD4075).

## Supplementary Figure

**Fig S1.**
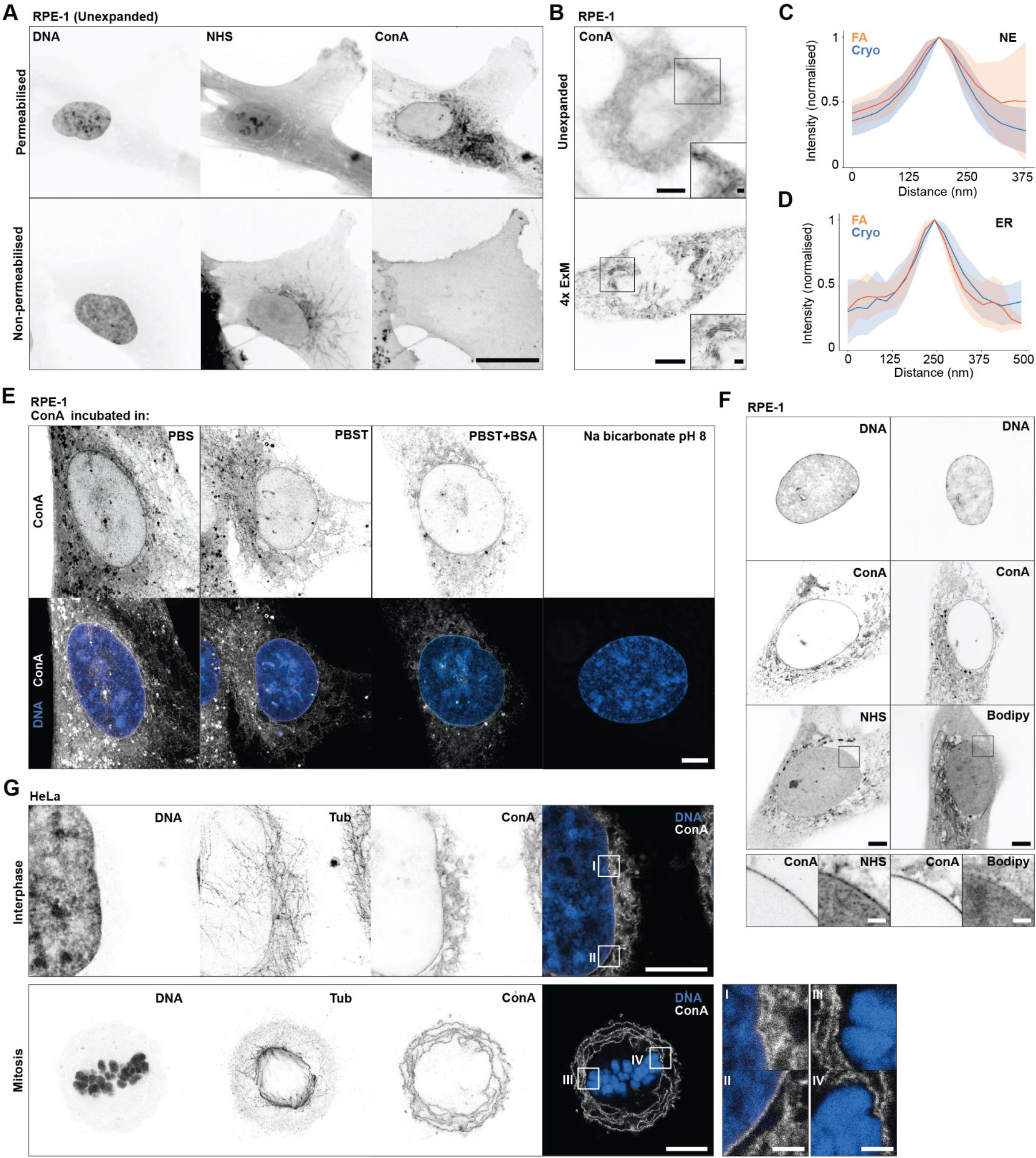
ConA staining conditions across unexpanded and expanded mammalian cells. A. Hoechst, NHS ester, and ConA staining of FA fixed, unexpanded RPE-1 cells either permeabilised with 0.1% Triton-X100 or left untreated (non-permeabilised). B. Comparison of ConA staining in unexpanded and expanded (4-fold U-ExM) mitotic RPE-1 cells. Squares represent zoom ins of membrane stacks. C. Quantification of ConA signal width at the NE in plunge frozen and FA-fixed RPE-1 cells. Distances are adjusted for the expansion factor. Line represents mean value, shaded areas represent standard deviation. N = 20. D. Quantification of ConA signal width at ER stacks in plunge frozen and FA-fixed RPE-1 cells. Distances are adjusted for the expansion factor. Line represents mean value, shaded areas represent standard deviation. N = 30. E. Comparison of ConA (grey) staining using PBS, PBS-T (0.2% Tween20), sodium bicarbonate pH8, or 3% BSA in PBS-T. DNA is shown in blue. Channel brightness is scaled the same across all conditions. F. Comparison of ConA labelling to pan-protein labelling (NHS ester) and pan-lipid labelling (Bodipy-Ceramide). Squares represent zoom ins of nuclear membrane. G. Hoechst, tubulin, and ConA staining of FA fixed, expanded HeLa cells during interphase and mitosis. DNA is shown in blue. Squares represent zoom ins of the nucleus. All scale bars are adjusted for the expansion factor and represent 5µm in overview images and 1µm in close ups.

**Fig S2.**
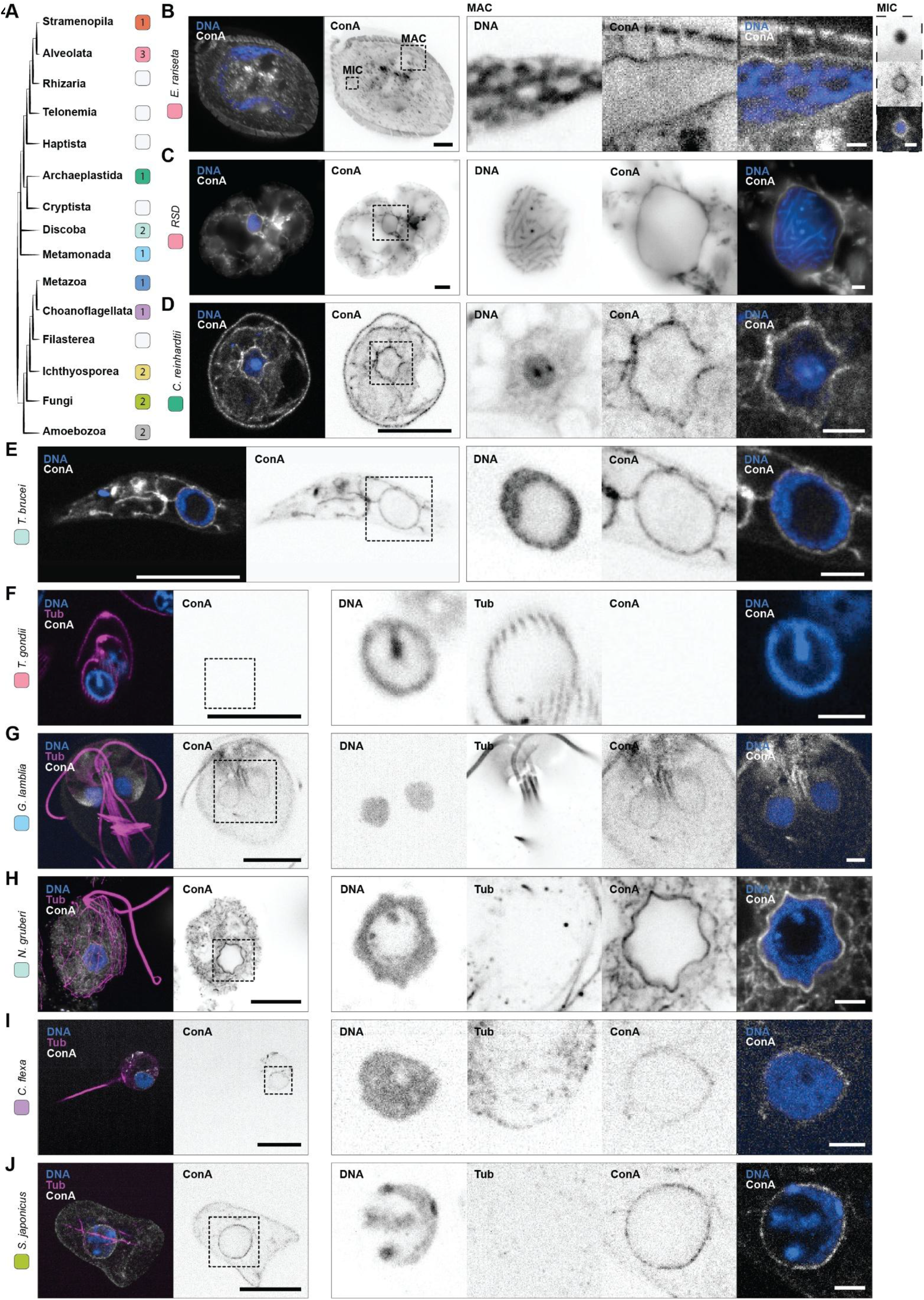
ConA staining across diverse eukaryotes. A. Cladogram of eukaryotes with colours indicating the position and number of used species within present figures. B. MIP image of the ciliate *Euplotes rariseta* (FA fixation). Crops show zoom-ins of the macronucleus (MAC) and micronucleus (MIC). C. Central slice images of Ross Sea dinoflagellate (FA fixation). Crop show zoom-in of the nucleus. D. Central slice images of green algae *Chlamydomonas reinhardtii*. Crop show zoom-in of the nucleus. E. MIP image of the kinetoplastid parasite *Trypanosoma brucei* (FA fixation. Crop show zoom-in of the nucleus and is a single slice. F. MIP image of the apicomplexan parasite *Toxoplasma gondii* (FA fixation. Crop show zoom-in of the nucleus and is a single slice. G. MIP image of *Giardia lamblia* (FA fixation). Crop show zoom-in of the nuclei and is a single slice. H. MIP image of *Naegleria gruberi* (FA fixation). Crop show zoom-in of the nucleus. I. MIP image of *Choanoeca flexa (*sequential fixation of 12% Acetone and 8% FA in 4xPBS). Crop show zoom-in of the nucleus and is a single slice. J. MIP image of the fission yeast *Schizosaccharomyces japonicus* (HPF fixation). Crop show zoom-in of the nucleus and is a single slice. For all merged images, ConA is in grey, DNA in blue, and tubulin, if shown, in magenta. All scale bars are adjusted for the expansion factor and represent 5µm in overview images and 1µm in close ups.

